# The lncRNA *Neat1* is associated with astrocyte reactivity and memory deficits in a mouse model of Alzheimer’s disease

**DOI:** 10.1101/2023.05.03.539260

**Authors:** Ashleigh B Irwin, Verdion Martina, Silvienne C Sint Jago, Rudhab Bahabry, Anna Maria Schreiber, Farah D. Lubin

## Abstract

Dysregulation of long non-coding RNAs (lncRNAs) have been associated with Alzheimer’s disease (AD). However, the functional role of lncRNAs in AD remains unclear. Here, we report a crucial role for the lncRNA *Neat1* in astrocyte dysfunction and memory deficits associated with AD. Transcriptomics analysis show abnormally high expression levels of *NEAT1* in the brains of AD patients relative to aged-matched healthy controls, with the most significantly elevated levels in glial cells. In a human transgenic APP-J20 (J20) mouse model of AD, RNA-fluorescent *in situ* hybridization characterization of *Neat1* expression in hippocampal astrocyte versus non-astrocyte cell populations revealed a significant increase in *Neat1* expression in astrocytes of male, but not female, mice. This corresponded with increased seizure susceptibility in J20 male mice. Interestingly, *Neat1* deficiency in the dCA1 in J20 male mice did not alter seizure threshold. Mechanistically, *Neat1* deficiency in the dorsal area CA1 of the hippocampus (dCA1) J20 male mice significantly improved hippocampus-dependent memory. *Neat1* deficiency also remarkably reduced astrocyte reactivity markers suggesting that *Neat1* overexpression is associated with astrocyte dysfunction induced by hAPP/Aβ in the J20 mice. Together, these findings indicate that abnormal *Neat1* overexpression may contribute to memory deficits in the J20 AD model not through altered neuronal activity, but through astrocyte dysfunction.

## Introduction

Alzheimer’s disease (AD) is a progressive neurodegenerative disorder characterized by neuronal hyperexcitability ^1, 2^,profound episodic memory impairments ^3^, and other cognitive deficits ^4^. Therefore, further exploration of the mechanisms underlying key characteristics of AD is necessary for the development of novel therapeutic approaches. Dysregulation of long non-coding RNA (lncRNA) expression has been shown to be strongly associated with cognitive dysfunction and neurodegenerative disorders ^5–8^. Numerous lncRNAs are aberrantly overexpressed in AD ^9–11^ and there is mounting evidence that transcriptional and post-transcriptional regulation by lncRNAs may contribute to AD pathogenesis ^12–14^.

Nuclear paraspeckle assembly transcript 1 *(NEAT1)* is a highly conserved lncRNA with altered expression in a number of neurodegenerative diseases ^15^. *NEAT1* exists in two isoforms, the short canonically polyadenylated *NEAT1_1,* and the longer *NEAT1_2*^16, 17^. While both *NEAT1_1* and *NEAT1_2* are present in paraspeckle subnuclear bodies, *NEAT1_2* is a critical component of paraspeckles associated with recruitment and sequestration of transcriptional machinery ^18–20^. While *NEAT1_2* is believed to have low expression in the healthy adult brain^17^, *NEAT1_1* is more widely expressed, including in the central nervous system, and can regulate gene activation independent of paraspeckles ^21^. Prior studies have identified a role for *Neat1* in regulation of critical memory-associated genes and in hippocampus-dependent memory formation^22^. Recent studies suggest that *NEAT1* is associated with endocytosis related gene regulation and uptake of amyloid beta oligomers by glial cells *in vitro* ^23^, as well as regulation of cytoskeletal components via interaction with microRNAs (miRNAs) in a mouse model of AD ^24^. Whether *Neat1* contributes to glia cell function and/or memory impairments in remains elusive.

The hippocampus is a complex, heterogeneous region necessary for proper memory formation and is particularly vulnerable to age-and disease-related damage ^25, 26^. The transgenic human-APP-J20 (J20) mouse model of AD develops gliosis and amyloid plaques in brain regions like the hippocampus contributing to hyperexcitability and memory deficits between four and six months of age^27^. By employing RNA-fluorescent *in situ* hybridization (FISH) to characterize expression of the highly conserved lncRNA *Neat1* in astrocyte and non-astrocyte cell populations in the hippocampus of the transgenic J20 mouse model of AD, with molecular and anatomical assessment, seizure behavior, and cognitive test, we demonstrate that *Neat1* contributes to astrocyte reactivity and memory deficits. We identified a significant increase in the expression of *Neat1* in astrocytes of male, but not female J20 mice that corresponded to a decreased seizure threshold in males. These findings suggest that *NEAT1* dysfunction in the hippocampus of AD patients may contribute to memory deficits associated with the disorder.

## Results

### Cell-type specific *NEAT1* overexpression in Alzheimer’s disease patients

In prior studies, we found that *Neat1* levels were abnormally increased in the hippocampus of normal aged mice compared to young adults^22^, raising the possibility that age-related disorders like Alzheimer’s Disease (AD) may present with similar *Neat1* changes. First, we sought to determine whether *NEAT1* expression is altered in the brain of patients with AD. Using scREAD that collates multiple single cell and single nuclei RNA sequencing datasets^28^, we performed a comprehensive analysis of the single-cell RNA-seq Database for AD. We found that the average fold change of *NEAT1* expression is increased in the entorhinal cortex (EC) and superior frontal gyrus (SFG) across all cell types in AD compared to age and sex match controls (**Figure 1A**). Interestingly, *NEAT1* expression levels are decreased in the prefrontal cortex (PFC) of AD patients compared to controls (**Figure 1A**) suggesting a unique region dependent function for *NEAT1* in AD. We then looked at the relative expression of *NEAT1* in each cell type compared to all other cell types in both healthy controls (**Figure 1B**) and AD patients (**Figure 1C**). In both healthy controls and AD patients, the general trend (increased or decreased relative expression) was the same for all three brain regions analyzed. Our analysis showed an increase in *NEAT1* expression levels in astrocytes and microglia relative to other cell types. We also found decreased *NEAT1* expression levels in excitatory neurons and oligodendrocyte precursor cells across all brain regions. Decreased expression if *NEAT1* in excitatory neurons is consistent with previous studies showing *NEAT1* decreases with neuronal firing^29^ and known hyperexcitability in the AD brain^30^. Together, these data demonstrate differential *NEAT1* expression patterns across the brain of AD patients that is cell type specific.

**Figure 1.**
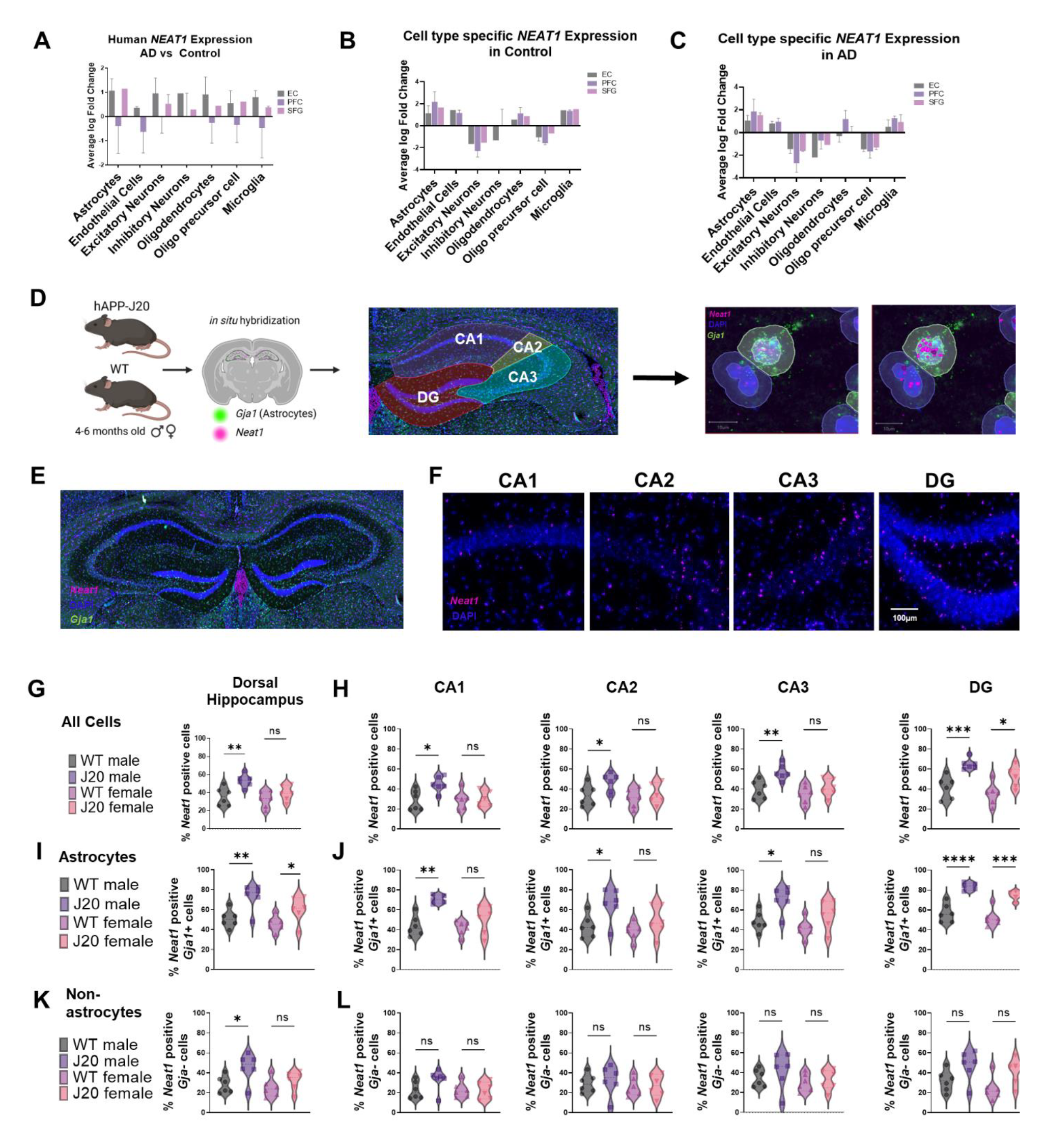
Increased glial expression of the lncRNA *NEAT1* in patients with AD and the male J20 mouse model of AD. A) Differential *NEAT1* expression from AD patients relative to age and sex-matched controls in three brain regions, Entorhinal Cortex (EC), Prefrontal Cortex (PFC) and Superior Frontal Gyrus (SFG). B) Average log Fold change of *NEAT1* by cell type relative to all other cell types in healthy controls. C) Average log Fold change of *NEAT1* by cell type relative to all other cell types in AD patients. Data from the scREAD Single-cell RNA-seq Database for Alzheimer’s Disease^28^ D) Schematic depicting experimental approach with an overlayed tracing of individual hippocampal subfields and representative image of *Neat1* positive cell detection in the J20 mouse model of AD. E) Representative RNAscope images taken on the Olympus slide scanner (40x) of dorsal hippocampus with *Neat1* (magenta) and DAPI staining. F) Representative images of each individual subfield (40x) G) Quantification of the proportion of *Neat1* positive cells in the dorsal hippocampus n=6 mice/group, One-way ANOVA followed by Tukey’s Multiple comparisons test *P<0.05, **P<0.01, ***P<0.001. H) Quantification of the proportion of *Neat1* positive cells in each subfield independently. n=6 mice/group, One-way ANOVA followed by Tukey’s Multiple comparisons test *P<0.05, **P<0.01, ***P<0.001. I) Quantification of the proportion of *Neat1* positive astrocytes (*Gja1+*) in the dorsal hippocampus. n=6 mice/group, One-way ANOVA followed by Tukey’s Multiple comparisons test *P<0.05, **P<0.01, ***P<0.001. J) Quantification of the proportion of *Neat1* positive astrocytes (*Gja1+*) in each subfield independently. n=6 mice/group, One-way ANOVA followed by Tukey’s Multiple comparisons test *P<0.05, **P<0.01, ***P<0.001. K) Quantification of the proportion of *Neat1* positive non-astrocytes (*Gja1-*) in the dorsal hippocampus n=6 mice/group, One-way ANOVA followed by Tukey’s Multiple comparisons test *P<0.05, **P<0.01, ***P<0.001. L) Quantification of the proportion of *Neat1* positive non-astrocytes (*Gja1-*) in each subfield independently. RNAscope data are n=6 mice/group, One-way ANOVA followed by Tukey’s Multiple comparisons test *P<0.05, **P<0.01, ***P<0.001.

### *Neat1* is overexpressed in the hippocampus of male, but not female, J20 mice

Next, we sought to determine if *Neat1* is altered in the brain of the J20 AD mouse model similar to AD patients. The hippocampus is a heterogenous structure with different regions responsible for different aspects of memory formation ^31^. Therefore, we bilaterally assessed *Neat1* expression in the Cornu Ammonis (CA1, CA2, CA3) and the Dentate Gyrus (DG) regions of the hippocampus. Each subfield was traced based on the Allen Mouse Brain Atlas^32^ and annotated in QuPath (**Figure 1D-F**). RNAscope probe analysis for *Neat1* puncta revealed that the proportion of cells expressing *Neat1* is significantly increased in male, but not female, J20 mice (**Figure 1G**). We found no significant difference in the average *Neat1* expression per cell between J20 and wild type (WT) littermates or between males and females (**Figure S1**). Interestingly, the proportion of cells expressing *Neat1* was found to be significantly increased in male J20 mice in CA1, CA2, and CA3, while there was an increase in *Neat1* expression in both male and female J20 mice in the DG (**Figure 1G**). Further quantification of individual *Neat1* puncta (**Figure S1D**) showed no significant differences between J20 and WT controls in either males or females. These findings demonstrate an increase in *Neat1* expression in the J20 mouse that recapitulates our findings in other brain regions critical for memory in AD patients.

### *Neat1* is increased in astrocytes, but not non-astrocyte cell populations in male J20 mice

Because we found significant changes in *NEAT1* expression levels specifically in glial cell populations in AD patients, we further investigated *Neat1* expression in astrocyte (*Gja1+*) versus non-astrocyte (*Gja1-*) cell populations in the hippocampus (**Figure 1I-L**). Using an RNAscope probe for the astrocyte marker *Gja1*, we found that at the level of the whole dorsal hippocampus, there is an increased proportion of astrocytes expressing *Neat1* in both male and female J20 AD mice (**Figure 1I**). However, within the non-astrocyte cell population, there were no significant differences between J20 mice and WT littermate controls, or between males and females (**Figure 1K**). Hippocampal subfield analysis revealed a significantly increased proportion of *Neat1* expressing cells in dorsal CA1, CA2, and CA3 in male J20 AD mice compared to controls, while both male and female J20 AD mice had an increased proportion of *Neat1* expressing cells in the DG (**Figure 1J**). No significant differences in the proportion of *Neat1* expressing cells were observed in any group within the non-astrocyte cell population (**Figure 1l**). We further investigated any differences in the average *Neat1* expression per *Neat1* positive cell and found no significant differences at the level of the whole dorsal hippocampus or within each hippocampal subfield within the astrocyte cell population (**Figure S1E, F**). However, we noted a trend towards a significant decrease in average *Neat1* expression levels in dorsal CA1 of female and male J20 AD mice (**Figure S1G**). Overall, our data suggest a shift in hippocampal astrocyte phenotype, as defined by expression of *Neat1* in the hippocampus of male J20 mice.

### Paraspeckles elements are not altered with total *Neat1* knockdown

While the long isoform *Neat1_2* and associated paraspeckles are not usually seen in the healthy brain^17^, paraspeckles are often increased in response to cellular stress^33^ associated with the inflammation seen in neurodegenerative diseases such as AD. Therefore, we sought to determine if manipulating *Neat1* in the J20 mouse model of AD would alter the expression of critical paraspeckle proteins (**Figure 2A**). Following siRNA mediated *Neat1* knockdown, Western blot analysis showed no difference in the levels of three key paraspeckle proteins, NONO, SFPQ, or FUS, in dCA1 of J20 animals compared to those who received a scrambled control (**Figure 2E-G**). Similarly, no changes in gene expression levels for these proteins were detected by qPCR (**Figure S2**). This suggests that transcriptional or functional impacts of increased *Neat1* expression in AD are not mediated by altered paraspeckle formation.

**Figure 2.**
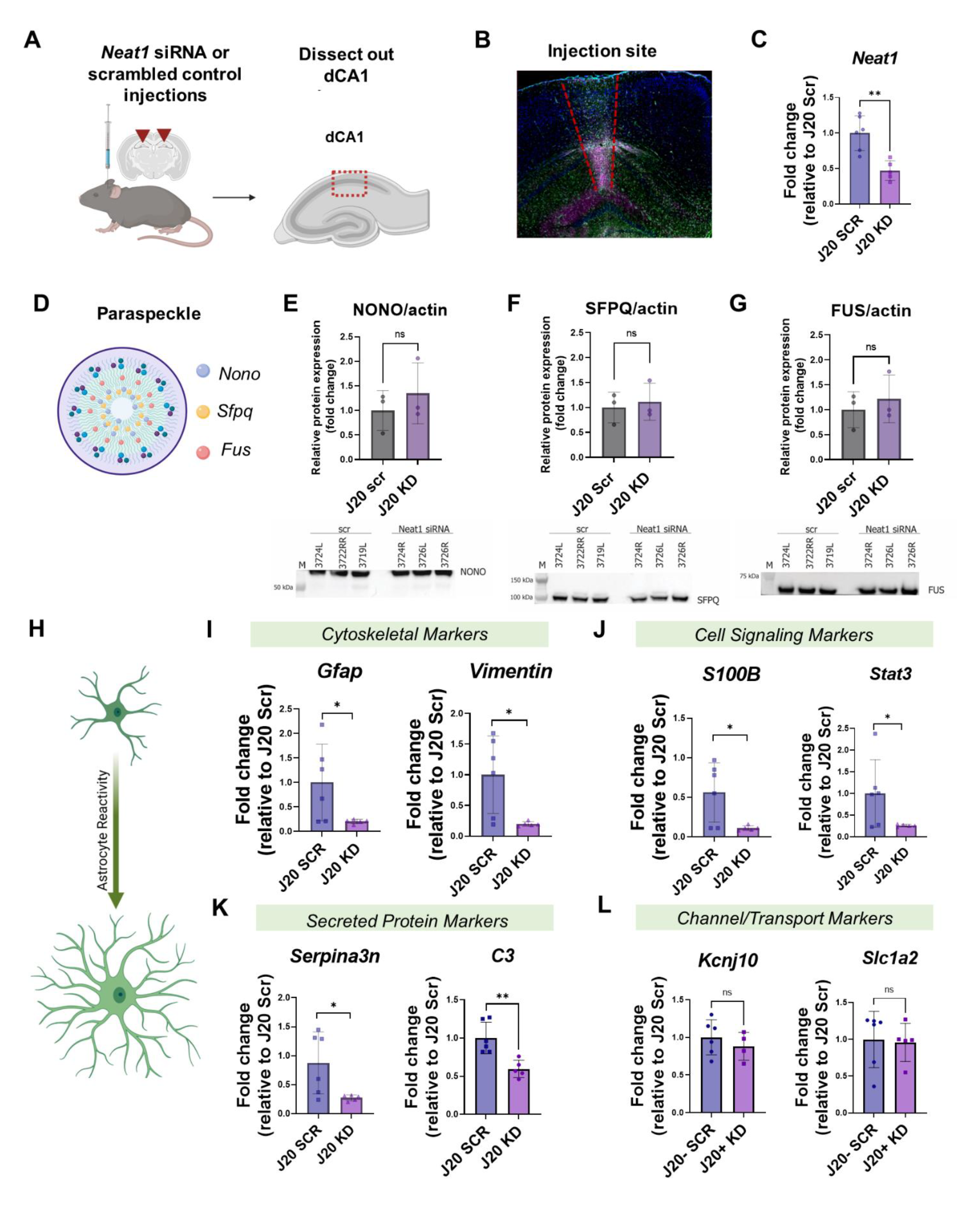
Hippocampal *Neat1* mediates astrocyte reactivity in the J20 mouse model of AD. A) Schematic of experimental design. B) Representative injection site image. C) Reverse transcription quantitative polymerase chain reaction (qPCR) showing *Neat1* knockdown relative to actin. Data are from n=3 mice/group Student’s t-test, ns = not significant*P<0.05, **P<0.01, ***P<0.001 D) Schematic of paraspeckle E-G) Western blot quantification of three key paraspeckle proteins, NONO, SFPQ, and FUS in *Neat1* siRNA treated J20 mouse as well as WT and J20 scrambled siRNA treated controls. Data are from n=3 mice/group Student’s t-test, ns = not significant H) Schematic of the transition from quiescent to reactive astrocyte I-L) qPCR quantification analysis of several molecular markers of astrocyte reactivity: *GFAP, Vimentin*, *S100B, Stat3*, *Serpina3n, C3,* and *Eaat1, Kcnj10.* Data are from n=5-6 mice/group (males and females) Student’s t-test, ns = not significant, *P<0.05, **P<0.01, ***P<0.001

### mRNA levels of genes associated with astrocyte reactivity are decreased with *Neat1* knockdown

With our J20 mice data recapitulating the glial predominance of *Neat1* expression changes seen in humans with AD, we sought to determine if *Neat1* expression impacted a key feature of astrocytes: astrocyte reactivity. Astrocyte reactivity is highly associated with AD^34^. As the primary output of the hippocampus and one of the first regions effected by AD pathology, we chose to focus on dCA1. To test the effect of reducing *Neat1* expression in male J20 mice, we directly infused *Neat1* targeting siRNAs into dCA1. Five days after intra-dCA1 injection of *Neat1* siRNA, we used qPCR analysis to quantify several gene markers of astrocyte reactivity. It is now recognized that astrocyte reactivity represents a complicated biological process that cannot be adequately quantified by simple *Gfap* expression^35^. Instead, reactivity analysis should encompass a variety of different cellular processes that are altered in reactive astrocytes. Therefore, we investigated gene expression levels of multiple cytoskeletal, cell signaling, secreted protein, and ion channel/transporter markers following *Neat1* knockdown. Significantly, we observed that gene expression of cytoskeletal (*Gfap, Vimentin*), cell signaling (*S100b, Stat3*), and secreted protein (*Serpina2n*) markers of astrocyte reactivity were all decreased in dCA1 with *Neat1* knockdown (**Figure 2 H-J**). We did not however observe any changes in channel/transport gene levels (*Kcnj10*, *Slc1a1*). This suggest increased *Neat1* in the J20 mouse model of AD contributes to AD-associated astrocyte reactivity, particularly in terms of altered cell signaling and immune signaling.

### *Neat1* knockdown in dCA1 does not impact seizure threshold in male J20 mice

A prominent feature of AD is the development of neuronal hyperexcitability, and decreased seizure threshold is well established in rodent models of AD^36–38^. Furthermore, neuronal hyperexcitability has been postulated as a potential mechanism driving the most notable consequence of AD, memory impairment^39^. Astrocytes have long been known as key homeostatic regulators of the brain, while increased astrocyte reactivity can lead to the progression of hyperexcitability and even the development of epilepsy^40–42^. Therefore, we sought to determine if *Neat1* expression in the J20 mice impacted seizure threshold. Towards this end, we used a Pentylenetetrazol (PTZ) challenge (45mg/kg) to first confirm an altered seizure threshold in the J20 mouse model of AD (**Figure 3A**). We found a decreased seizure threshold in male but not female J20 mice using a PTZ challenge (**Figure 3B,C**). To determine if *Neat1* knockdown would impact seizure threshold, male J20 mice and WT littermates received intra-dCA1 infusions of *Neat1* siRNA or scrambled control siRNA (**Figure 3D**). Five days following the infusion of siRNA, we found no significant differences in max score, time to forelimb clonus or time to max stage in response to 45mg/kg IP PTZ (**Figure 3E**). While behavioral status following PTZ administration was not impacted by *Neat1* knockdown, we also sought to determine if *Neat1* knockdown impacted neuronal activity at subthreshold levels. qPCR for *Fosb* was used as a marker for chronic neuronal activity and showed no change with *Neat1* manipulation after PTZ challenge (**Figure 3F**). Together this suggests that increased *Neat1* in dCA1 does not contribute to decreased seizure threshold or hyperexcitability in the J20 mouse model of AD. Interestingly, gene markers for astrocyte reactivity were found to be decreased with *Neat1* knockdown following PTZ challenge. While astrocyte reactivity is associated with altered neuronal excitability, our data suggests that *Neat1* mediated reactivity is not necessary for decreased seizure threshold in the J20 mouse model of AD.

**Figure 3.**
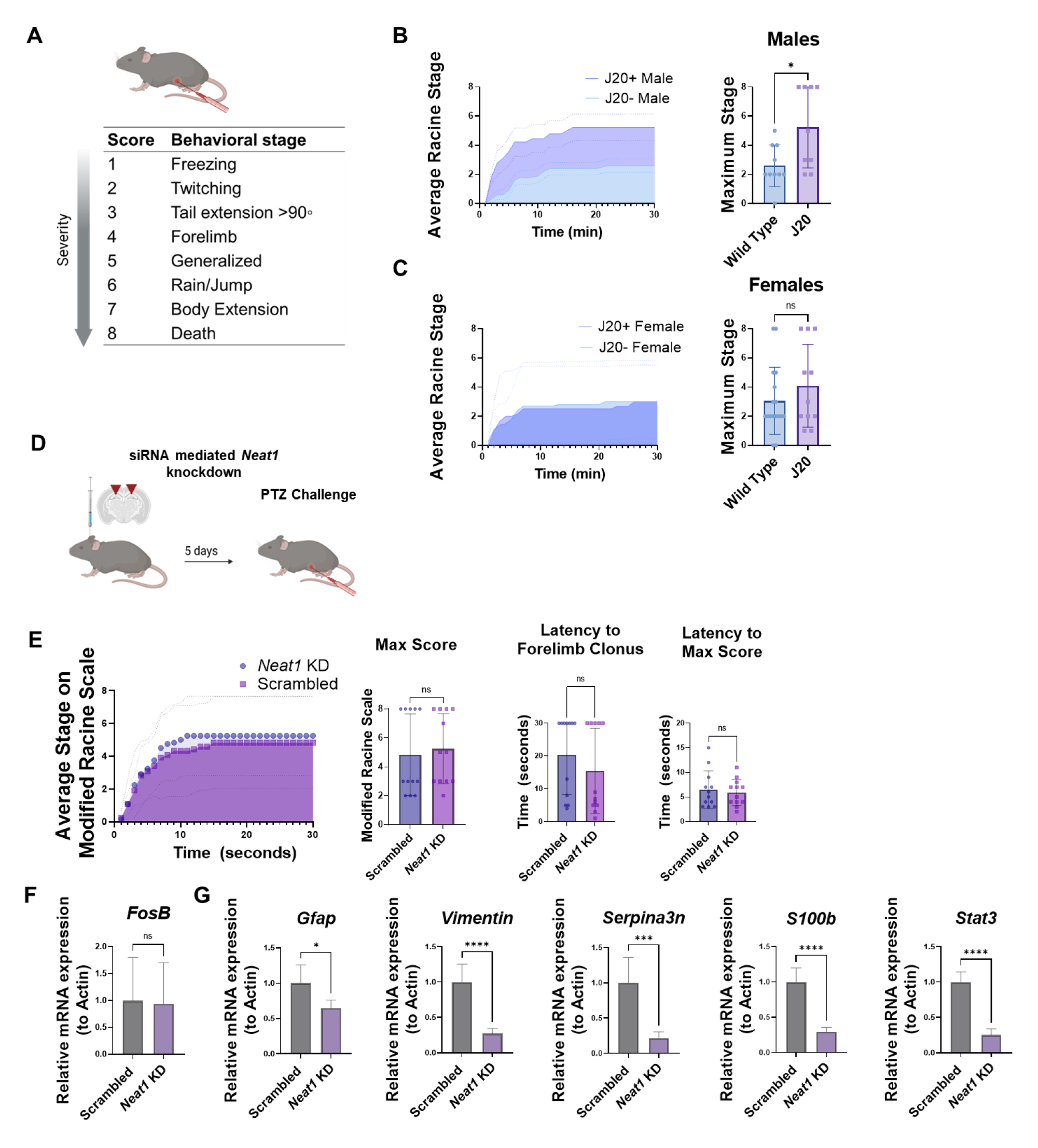
siRNA mediated *Neat1* knockdown in dCA1 does not alter seizure threshold in the J20 mouse model of AD. A) Graphical depiction of the modified Racine scale used to quantify seizure stage following Pentylenetetrazol (PTZ) injection (45mg/kg). B) Left, Quantification of average Racine stage by minute over the course of 30 minutes following PTZ injection in J20 males and wild type (WT) littermate controls. Right, average maximum stage reached. Data is from n=9-10 male mice/group Student’s t-test *P<0.05, **P<0.01, ***P<0.001 C) Left, Quantification of average Racine stage by minute over the course of 30 minutes following PTZ injection in J20 females and WT littermate controls. Right, average maximum stage reached. Data is from n=13-17 female mice/group. Student’s t-test *P<0.05, **P<0.01, ***P<0.001 D) Graphical depiction of siRNA mediated *Neat1* knockdown or scrambled siRNA control injections in dCA1 of male J20 mice followed five days later by PTZ challenge and subsequent qPCR of injection site. E) Quantification of average Racine stage over 30 minutes, Maximum Racine stage, Latency to Forelimb Clonus and Latency to maximum Racine stage. F-G) qPCR quantification of *Fosb* and the astrocyte reactivity markers *GFAP, Vimentin*, *S100B, Stat3*, *Serpina3n*. Data from n=12 male mice/group. Student’s t-test *P<0.05, **P<0.01, ***P<0.001

### *Neat1* knockdown in dCA1 improves hippocampus-dependent memory

Area CA1 of the hippocampus is important for associative long-term memory and becomes dysfunctional with normal aging and age-related disorders such as AD^43, 44^. We next sought to determine the effect of decreasing *Neat1* expression levels on hippocampus-dependent memory. Male and female J20 AD mice and WT littermates received intra-dCA1 infusions of *Neat1* siRNA or scrambled control siRNA (**Figure 4A**). Five days following siRNA infusion, we confirmed decreased *Neat1* expression levels in dorsal CA1 (**Figure 4B**). In a separate cohort of animals, we performed a series of hippocampus-dependent behavioral tasks, including open field habituation, context fear conditioning, and Y maze spontaneous alternation test, to assess long-term memory and working memory, respectively. No significant differences in anxiety behavior as measured by thigmotaxis were observed using an open field apparatus (**Figure 4C**). To assess working memory, animals were tested in the Y maze behavior task, and spontaneous alternations were recorded. We observed no differences in the frequency of alternations or the maximum number of alternations between any of the groups (**Figure 4D, E**). This suggests that 4- to 6- month-old J20 mice lack deficits in working memory and *Neat1* knockdown does not impact their performance in the Y maze behavior task. However, J20 mice with *Neat1* knockdown in dCA1 showed a significant decrease in cumulative distance traveled 24 hours after training in an open field habituation task (**Figure 4F, G**). Open field habituation exploits the innate tendency of mice to explore a novel environment, and the significantly decreased exploration on day two, suggests improved memory of the context relative to J20 mice who receive scrambled control injections. When tested in a context fear conditioning paradigm, there was no significant difference in freezing behavior between the three groups before or after foot shock. However, 24 hours later, when the animals were returned to the context, we observed increased freezing in J20 mice with *Neat1* knockdown relative to control J20 mice, and at levels similar to those seen in WT controls (**Figure 4H**). Similar to the open field habituation paradigm, this data suggests improved hippocampus-dependent memory in J20 mice with *Neat1* knockdown. We observed no significant differences in distance traveled or average velocity between groups (**Figure S3A**) indicating that the differences detected in memory behavior are not driven by altered activity levels, either in the J20 mice compared to WT controls, or with *Neat1* knockdown. Together, these findings suggest that *Neat1* knockdown in the dorsal hippocampus is sufficient to improve hippocampus-dependent memory formation in the J20 mouse model of AD.

**Figure 4.**
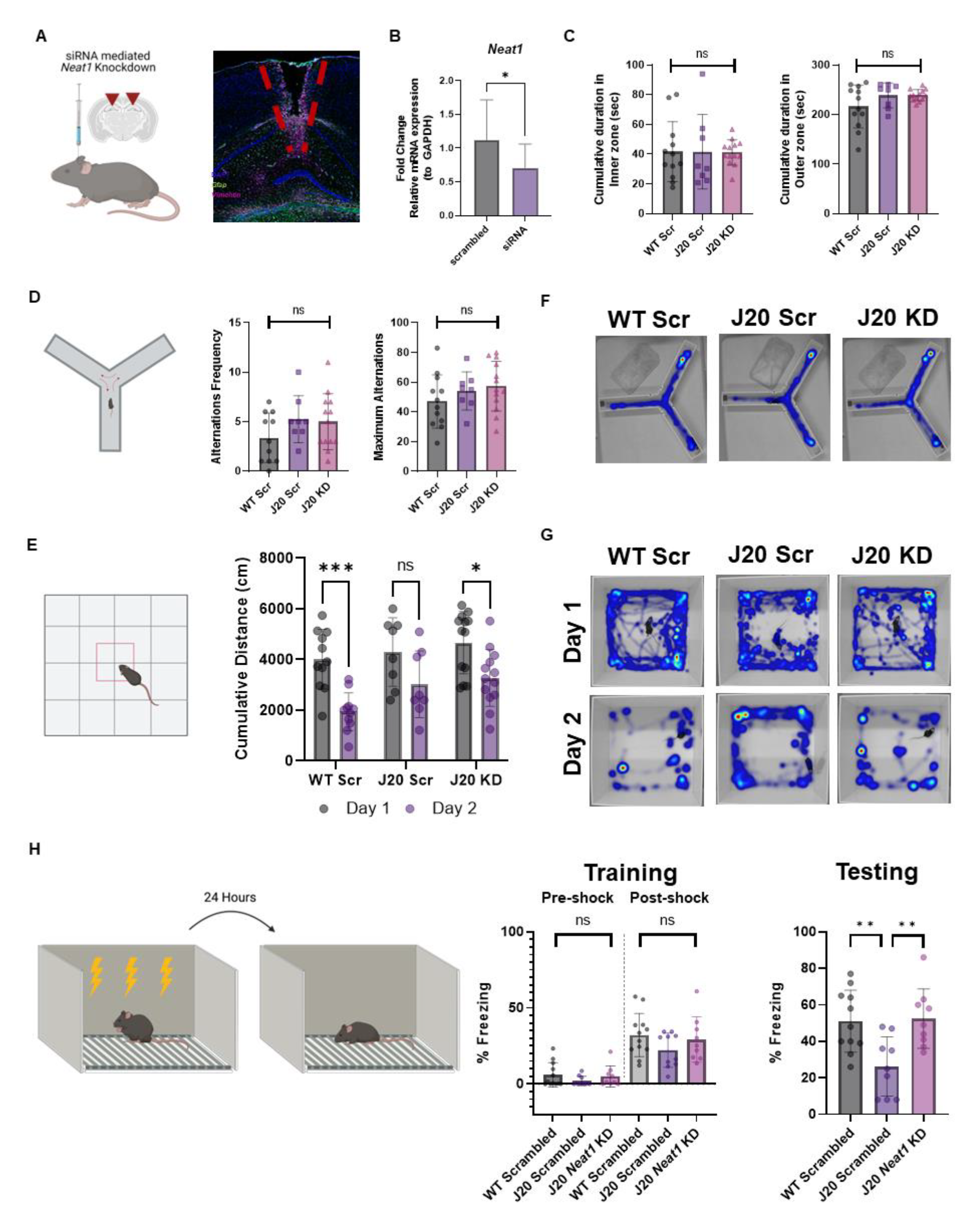
*Neat1* mediates hippocampus-dependent long-term memory in the J20 mice. A) Graphical depiction of siRNA mediated *Neat1* knockdown in dCA1 and representative injection site image. B) qPCR quantification of *Neat1* knockdown five days following dCA1 injection. N=4/group, mice were 4-6 months of age. Student’s t-test *p<0.05. C) Quantification of thigmotaxis following dCA1 *Neat1* Knockdown based on (left) cumulated duration in the inner zone (seconds) and (right) cumulative duration in the outer zone (seconds). Data is n=8-13 male and female mice One-way ANOVA ns = not significant, p>0.05. D) Graphical depiction (left) and quantification of alternation frequency (middle) and Maximum Alternations in a Y maze following siRNA knockdown in dCA1 of the J20 mouse model of AD. Data is n=8-13 male and female mice. One-way ANOVA followed by Tukey’s multiple comparisons test ns = not significant. E) Representative Y maze heatmaps from each group. F) Graphical depiction of Open field habituation task (left) and quantification of cumulative distance traveled (cm) on day 1 of training and 24 hours following. Data is n=8-13 male and female mice. Two-way ANOVA followed by Bonferroni’s comparisons test ns = not significant, p>0.05. *p<0.05. **p<0.01. G) Representative open field heatmaps from each group. H) Graphical depiction of contextual fear conditioning paradigm five days following bilateral infusion of *neat1* targeting siRNA or scrambled siRNA control (left). Freezing behavior was quantified before and after shock (middle) and again 24 hours later when animals were returned to the context (right). Data are n = 10-12 male and female mice/group. mice were 4-6 months of age. One-way ANOVA followed by Tukey’s multiple comparisons test *p<0.05. **p<0.01

## Discussion

While several studies have revealed significant dysregulation of lncRNAs in AD ^11, 14^ and explored their potential use as biomarkers ^45–47^, our understanding of the functional consequences of altered lncRNA expression levels in AD remains unexplored. *Neat1* is a highly conserved lncRNA that plays a critical role in several biological functions ^21, 48^; many of these functions are implicated in AD, including uptake of amyloid^23^, regulation of neuronal activity^29^ and memory consolidation^22^. To our knowledge, our data show for the first time that hippocampal *Neat1* is increased in astrocyte populations in J20 mice, and that this increase contributes to astrocyte reactivity and to associated memory impairment in the J20 AD mouse model.

In this study we demonstrate a sex and cell-type specific difference in *Neat1* expression the dCA1 of the hippocampus, with significantly increased astrocytic *Neat1* in male J20 animals relative to controls. This is significant considering the well-established sexual dimorphisms in the prevalence and progression of AD^49, 50^. Elucidation of the sex specific mechanisms driving AD pathology has considerable implications for the development of improved therapeutic strategies and personalized medicine. In addition, we saw a corresponding sex difference in seizure threshold, with male J20, but not female J20 mice, showing reduced seizure threshold (**Figure 3B, C**). This is consistent with studies that have shown a protective effect of a second X chromosome on mortality and cognitive impairment, particularly in the J20 mouse model of AD^51^. Astrocyte-mediated neuronal hyperexcitability is well described in both humans with AD^30^ and rodent models of AD^40, 52^. Therefore, we hypothesized that *Neat1* contributes to abnormal neural excitability in the J20 AD mice. While our *Neat1* manipulation in dCA1 did not impact seizure threshold, *Neat1* may still play a role in associated excitability at subthreshold levels, or in other hippocampal areas in the J20 mice.

The predominance of *Neat1* changes in the astrocyte cell population led us to hypothesize that increased *Neat1* expression was driving altered astrocyte functionality. We demonstrate that siRNA knockdown of *Neat1* in dCA1 results in decreased expression of multiple, diverse cellular markers of astrocyte reactivity. The transition to what has classically been described as astrocyte reactivity, is a molecular and functional transformation highly associated with neurodegenerative diseases ^34, 53, 54^. Moreover, decreasing astrocyte reactivity has been shown to improve cognitive deficits in a mouse model AD ^55^. This data suggests that increased hippocampal *Neat1* in AD drives altered astrocyte function associated with reactive astrogliosis, and contributes to altered astrocyte gene expression profiles described in AD ^56^.

The most well characterized function of *Neat1* is in the assembly of subnuclear paraspeckle bodies. Paraspeckles are subcellular bodies seen in response to cellular stress ^19^, and are capable of mediating significant changes in transcription regulation ^20^. Though paraspeckles are not generally believed to be present in high expression levels *in vivo* in the healthy brain, they are upregulated in response to stress and have been proposed to play a role in neurodegenerative diseases^15, 33^. We observed no changes in gene or protein expression of the critical paraspeckle components *Nono*, *Sfpq* or *Fus* between J20 animals and age matched WT littermates, nor any differences in expression upon siRNA knockdown of *Neat1*. These results indicate that altered transcriptional regulation due to increased *Neat1* may be mediated independent of paraspeckle bodies.

Recent studies show that decreases in *Neat1* expression following learning is necessary for memory consolidation^22^. Therefore, we hypothesized that *Neat1* contributes to AD-related cognitive impairments. Indeed, we saw significant improvement in hippocampus-dependent memory with *Neat1* knockdown in dCA1 of the J20 mouse model of AD. While *Neat1* is known to mediate memory during normal aging, the J20 mouse model of AD develops key features of AD, including gliosis, hyperexcitability and memory deficits by four to six months of age^27^, suggesting that the contribution of increased *Neat1* to AD-related memory impairment are independent of age. Neuronal hyperactivity has been proposed as a contributor to the progression of memory dysfunction in AD ^39, 57^, yet lack of impact on seizure threshold following *Neat1* knockdown indicates *Neat1* contributes to cognitive function independently of neuronal hyperexcitability.

Overall, these findings indicate that *Neat1* mediates cognitive deficits through associated insult such as altered astrocyte function in the J20 mice, providing insight into the elusive role lncRNA and astrocytes play in the pathogenesis of AD.

### Limitations of the study

There are several limitations to this study. In the current study, we used the *Gja1+/Gja1*- label to distinguish between astrocyte-specific populations and other cell types. While the *Gja1+* marker likely encompass the majority of hippocampal astrocytes, we acknowledge that this approach limits our study to the analysis of *Gja1+* astrocytes, which may exclude smaller populations of *Gja1-* astrocytes. With the increased knowledge of astrocyte diversity ^56, 58, 59^ further studies should address the role of *Neat1* in the context of astrocyte heterogeneity. Indeed, Grubman et al., described 8 distinct populations of astrocytes, with a shift to two predominant subgroups in AD^56^. The *Neat1* knockdown approach utilizing siRNA results in pancellular *Neat1* knockdown, which does not consider astrocyte heterogeneity, which underlies astrocyte functional diversity. Future studies should consider cell-type-specific targeting methods to elucidate the impact of reducing *Neat1* specifically in astrocytes in the context of AD.

## Methods

### Resource availability

#### Lead Contact

Further information and requests for resources should be directed to and will be fulfilled by the lead contact, Farah D. Lubin

#### Materials availability

This study did not generate new unique reagents.

### Human gene expression and analysis

Data from three human AD single cell RNA sequencing datasets was collected using the scREAD Single-cell RNA-seq Database for Alzheimer’s Disease^28^. To determine differential *NEAT1* expression between AD patients and healthy individuals, data sets were filtered within the scREAD database for AD patients compared to age-and sex-matched controls within individual brain regions (Entorhinal cortex, Prefrontal cortex and Superior frontal gyrus). Differential expression values were averaged and plotted as the average log-fold change in gene expression relative to controls. Alternatively, to determine differential gene expression by cell type, data sets were filtered within scREAD by cell type and either control or AD, independently. Values are plotted as the average log-fold change in expression relative to the average *NEAT1* expression of all other cell types.

### Experimental Model and subject details

#### Animals

Male and female transgenic human APP J20 mice as well as transgene-negative littermate controls between four and six months of age were used for this study. Mice were group-housed (two to seven animals/cage) in plastic cages with *ad libitum* access to food and water and maintained on a 12-hour light/dark cycle. Behavioral tests were performed during the light cycle and approved by the University of Alabama at Birmingham Institutional Animal Care and Use Committee in and conducted in accordance with the National Institute of Health ethical guidelines.

#### Whole brain collection and sectioning

Animals were anesthetized using isoflurane and transcardially perfused with ice-cold 1x Phosphate Buffered Saline (PBS). Whole brains were removed and immediately placed on dry ice for later sectioning. Fresh frozen brains were then embedded in O.C.T and serial coronal sections (14 µm) were collected using a Leica CM1950 cryostat. Slides were stored at −80°C prior to immunofluorescence or *in situ* hybridization.

#### RNAscope

To characterize expression on *Neat1* throughout the hippocampus, RNAscope was performed on fresh frozen sections on slides (14 µm) using RNAscope Multiplex Fluorescent Reagent Kit V2 according to manufacturer’s recommendations.

Briefly, slides from frozen brains were transferred directly from −80°C to pre-chilled 10% normal buffered formalin and incubated for 15 mins at 4 °C before rinsing in 1x PBS. Slides were then dehydrated sequentially in 50%, 70% and 100% ethanol for 5 minutes at room temperature (RT). Slides were allowed to air dry for 5 minutes and then each section was traced using a hydrophobic pen. Endogenous peroxidases were quenched using hydrogen peroxide for 10 minutes at RT and antigen retrieval was performed using RNAscope Protease IV for 30 minutes at RT. RNAscope probes were warmed and C2 probe (*neat1*) was diluted 1:50 in C1 probe (*Gja1*). Approximately 100 µl of probe was added to each section and incubated in a humidity control tray for 2 hours at 40 °C in the Biotechne HybEZ oven. Slides were then washed in 1x Wash buffer (ACDbio) at room temperature and stored overnight in 5x Saline-sodium citrate.

Horse radish peroxidase signal was developed, and probes were detected with Opal dye. Slides were coverslipped using Invitrogen ProLong Diamond Antifade Mountant with DAPI and allowed to dry for two hours to overnight before imaging.

#### Imaging and quantification

RNAscope images were collected using an Olympus VS200 Research Slide Scanner. Both hemispheres of the hippocampus were selected, and z stacks of optical sections were collected at 40x and collapsed to a single image for quantification. Collapsed stacks were imported into QuPath software for analysis. Briefly, each subfield of the hippocampus (CA1, CA2, CA3 and DG) was traced bilaterally and annotated as the appropriate region. Three to four sections were traced per animal, with six animals per group. To identify the proportion of cells expressing *neat1*, cells were identified as astrocytes or other using a DAPI and *Gja1* (FITC) threshold in the Positive Cell Detection function of QuPath. Cells were then identified *neat1* positive or negative threshold (Cy3). The number of estimated *neat1* puncta was quantified using the subcellular detection function. Thresholds were set and visually confirmed for accuracy for each image.

#### siRNA delivery

Male and female mice between four and six months of age were anesthetized with isoflurane and received bilateral injections into dorsal CA1 (dCA1) of either Lincode SMARTpool siRNA (Dharmacon) targeting *neat1* or scrambled siRNA control. Injections were at stereotaxic coordinates AP −2.0mm, ML ± 1.5mm, DV −1.7mm with respect to bregma. siRNA stocks of 100 µM were diluted to ∼1.5 µM and conjugated with in vivo JetPei transfection reagent (Polyplus Transfection) on the day of surgery. Infusions were given at a rate of 0.1 µL per minute for a total volume of 1 µL per hemisphere. Animals were allowed to recover on a heating pad before being returned to their home cage. Behavioral studies (memory test or PTZ challenge) were performed beginning five days post-surgery.

#### Animal Behavior Assessment

Animals were transported to the behavior room on the morning of testing and allowed to habitat for a minimum of 30 minutes prior to testing. Two cohorts of animals were used to assess locomotion, anxiety, and memory behavior. One cohort was used to test thigmotaxis in the open field, open field habituation and spontaneous alternations in Y-maze. Due to the stressful nature of Context Fear Conditioning (CFC), a second cohort was used separately to test hippocampus-dependent memory in CFC.

#### Open Field Locomotor Test

Exploratory and anxiety-like behaviors were investigated using an open field apparatus (45cm x 45cm) and animal movements (distance traveled, velocity, duration in inner and outer zones) were recorded using Ethovision software. Mice were placed in the center of the open-field apparatus and allowed to explore freely for 5 minutes. The open field was divided into a grid with the inner zone composed of the inner half (four quadrants) and the outer zone defined as the wall-adjacent grid.

#### Open Field Habituation

To investigate long-term hippocampus-dependent memory, animals were tested twice in an open field apparatus described above, with the second trial occurring approximately 24 hours after the first trial. Cumulative distance traveled and average velocity were calculated across the two trials.

#### Y Maze

Working spatial memory was assessed in the animals by measuring the number of spontaneous alternations in a Y maze test. The maze consisted of three arms of equal length in a Y shape joined by a center region. Mice were placed in the center of the maze and allowed to explore freely for 5 minutes. The cumulative distance, velocity, number of arm entries and total number of alternations was recorded and calculated using the Noldus Ethovision software.

#### Contextual Fear Conditioning

Mice were trained in a contextual fear condition (CFC) paradigm in a novel context and then returned to the context 24 hours later, and freezing behavior was recorded by Med Associates software to assess long-term memory. The CFC paradigm consisted of a 119-second baseline period during which the animals were free to explore the context, followed by three shock pairings (0.5mA, 1s) with 59-second rest periods between each shock. Twenty-four hours later, the animals were returned to the context for five minutes to assess memory retention.

#### dCA1 Dissection

To assess gene expression and quantify protein levels, dorsal CA1 was collected around the site of siRNA/control injection. Animals were anesthetized using isoflurane and transcardially perfused with ice-cold PBS. Whole brains were removed and immediately placed on dry ice. dCA1 at the injection site was then dissected out with the aid of a mouse brain matrix (Harvard Apparatus) and tissue stored at −80°C prior to processing.

#### Isolation of protein and western blot

Samples were homogenized using lysis buffer containing 0.1 M HEPES, 1M magnesium chloride (MgCl_2_), 1M potassium chloride (KCl), 1M ditiotreitol (DTT), 10% Nonoxynol-40 (NP- 40), supplemented with phosphatase inhibitor (Sigma-Aldrich) and protease inhibitor (Sigma-Aldrich) cocktails. Nuclear extraction was performed using an extraction buffer containing 0.1 M HEPES, 1 M MgCl_2_, 1M DTT, 5 M sodium chloride (NaCl), 0.5 M ethylenediaminetetraacetic acid (EDTA), 50% glycerol, supplemented with phosphatase inhibitor (Sigma-Aldrich) and protease inhibitor (Sigma-Aldrich) cocktails. The concentration of isolated protein was measured using Quick Start™ Bradford Protein Assay (Bio-Rad).

Protein electrophoresis was performed on NuPAGE 4–12% Bis-Tris gels (Life Technologies) using thirty-five micrograms of samples, followed by transfer onto nitrocellulose membranes (Thermo Fisher). The following primary antibodies with respective dilutions were used for target detection: 1:10 000 anti-SFPQ (Abcam), 1:5000 anti-FUS (ProteinTech), 1:10 000 anti-NONO (Sigma-Aldrich), 1:10 000 anti-GFAP (Abcam), 1:100 anti-P-STAT3 (Cell Signaling), 1:1000 anti-Vimentin (Cell Signaling), 1:10 000 actin (Cell Signaling). For primary antibody detection, all membranes were incubated with 1: 7500 anti-rabbit IgG conjugated to horseradish peroxidise (HRP) (Thermo Fisher). Chemiluminescence of protein signals was detected using Clarity Western ECL Substrate (Bio-Rad) and quantified with ImageLab software (Bio-Rad).

#### qPCR

RNA was extracted using the Qiagen AllPrep DNA/RNA/Protein Mini kit as according to manufacturer’s instructions. Following extraction ∼100ng of RNA was DNAse treated (Amplification grade DNAse I, Sigma) and converted to cDNA (iScript cDNA synthesis kit; Biorad) and PCR amplified on the CFX1000 real-time PCR system (Biorad). Full descriptions of Thermo Scientific Taqman assays are in the supplemental materials. All datasets were analyzed using the delta-delta Ct method.

#### Statistical Analysis

Data from RNAscope and qPCR experiments were analyzed using one-way analysis of variance (ANOVA) followed by a Tukey’s post hoc test for multiple comparison or with Student’s t test. Behavior data were analyzed using the same methods apart from the open field habituation data which utilized a Two-way ANOVA followed by a Bonferroni’s post hoc analysis. Tests were two-tailed unless otherwise noted in the text. One-tailed tests were considered appropriate where changes in a particular direction were expected based on prior data. Outliers were identified in datasets using Grubb’s test (α =0.05) and any outliers excluded prior to analysis. Statistical tests were performed using GraphPad Prism 9. In all cases *n* represents the number of biological replicates or animals used (both collected tissue and in behavioral analysis).

## Acknowledgements and Funding

This work was supported by National Institute of Health (NIH) grants AG071785 (F.D.L.), and the Evelyn F. McKnight Brain Institute at the University of Alabama at Birmingham (F.D.L.). V.M., R.B., and S.S.J. were supported in part by the NIH/NINDS graduate Neuroscience Roadmap scholar training grant R25NS089463. A.I. is supported by the NIH/NINDS T32 NS061788. The Olympus slide scanner used in these studies was supported by the P20AG068024 grant.

## Author Contributions

Conceptualization, F.D.L. and A.B.I.; Methodology, F.D.L. and A.B.I; Investigation, A.B.I., V.M, S.S.J., R.B., A.S., and F.D.L.; Visualization, A.B.I.; Writing – Original Draft, A.B.I. and F.D.L.; Writing – Review & Editing, F.D.L., A.B.I., V.M., S.S.J., R.B., A.S., Funding Acquisition, F.D.L.

## Competing Interests

All authors declare no competing interests.

## Materials & Correspondence

All materials and correspondence related to data used in this paper should be addressed to Dr. Farah D. Lubin.

## Notes

### Competing Interest Statement

The authors have declared no competing interest.

