## Supplementary figures and images for "The lncRNA *Neat1* is associated with astrocyte reactivity and memory deficits in a mouse model of Alzheimer’s disease"

### Supplemental Figure 1

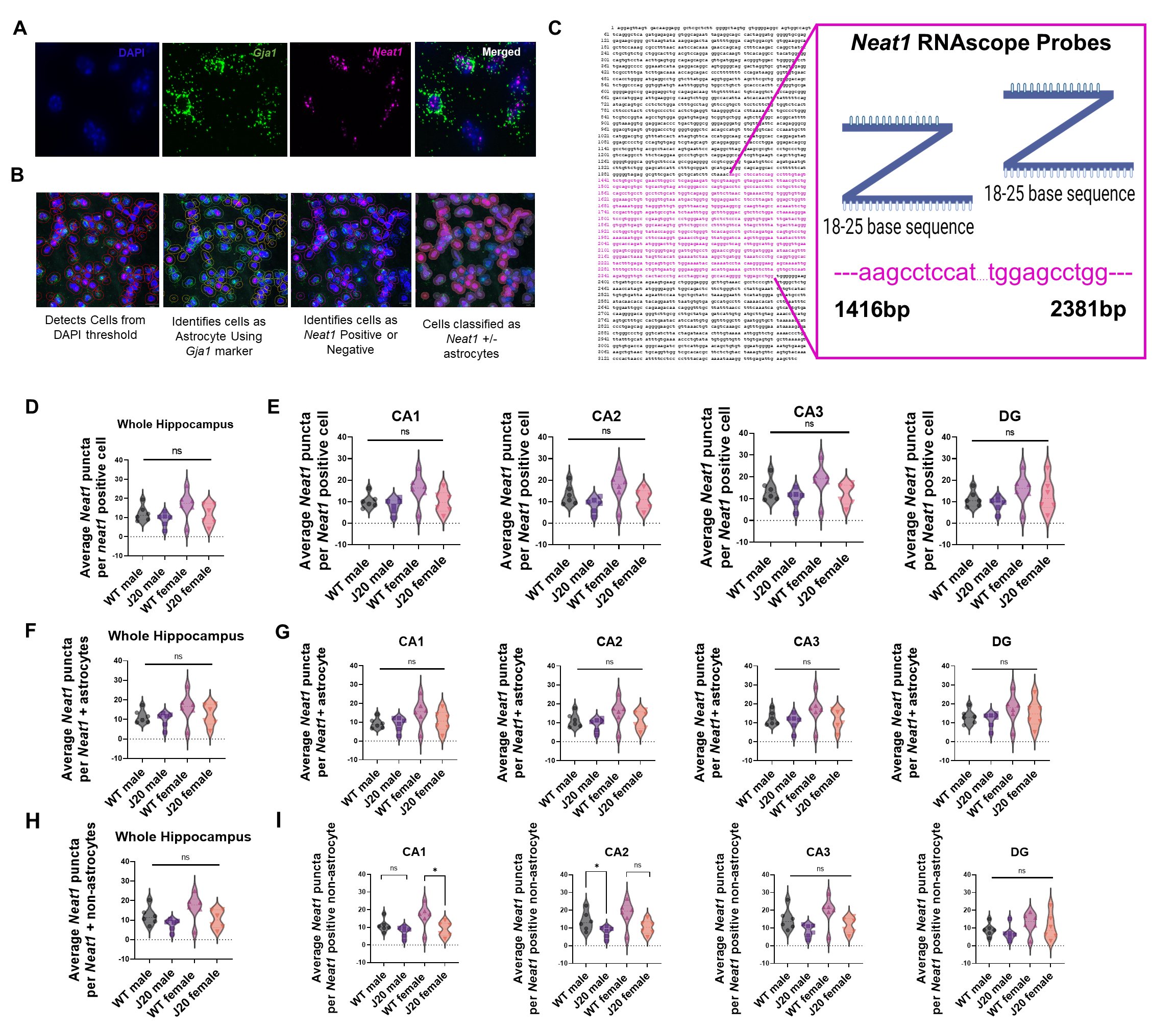

### Supplemental Figure 2

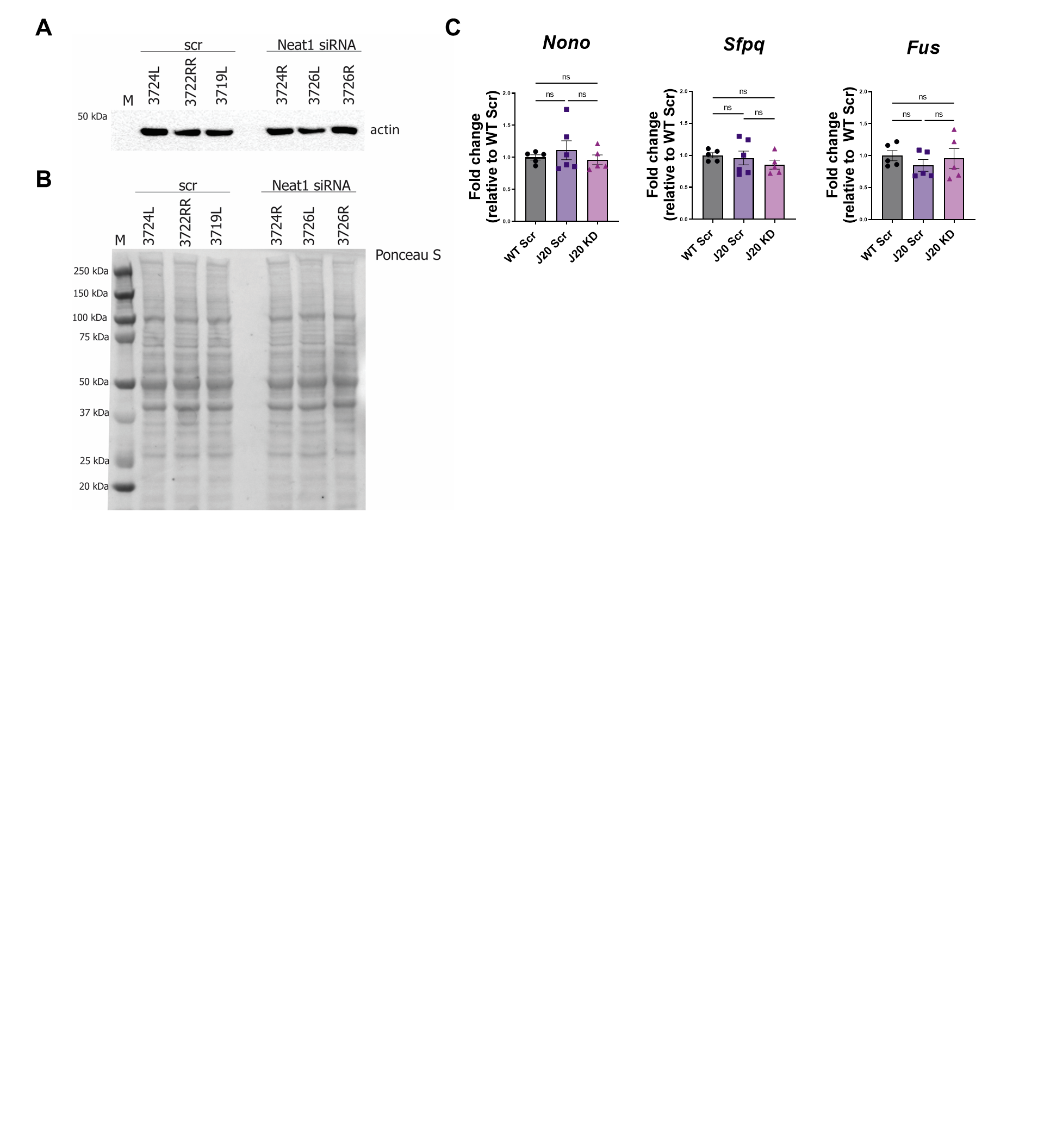

### Supplemental Figure 3

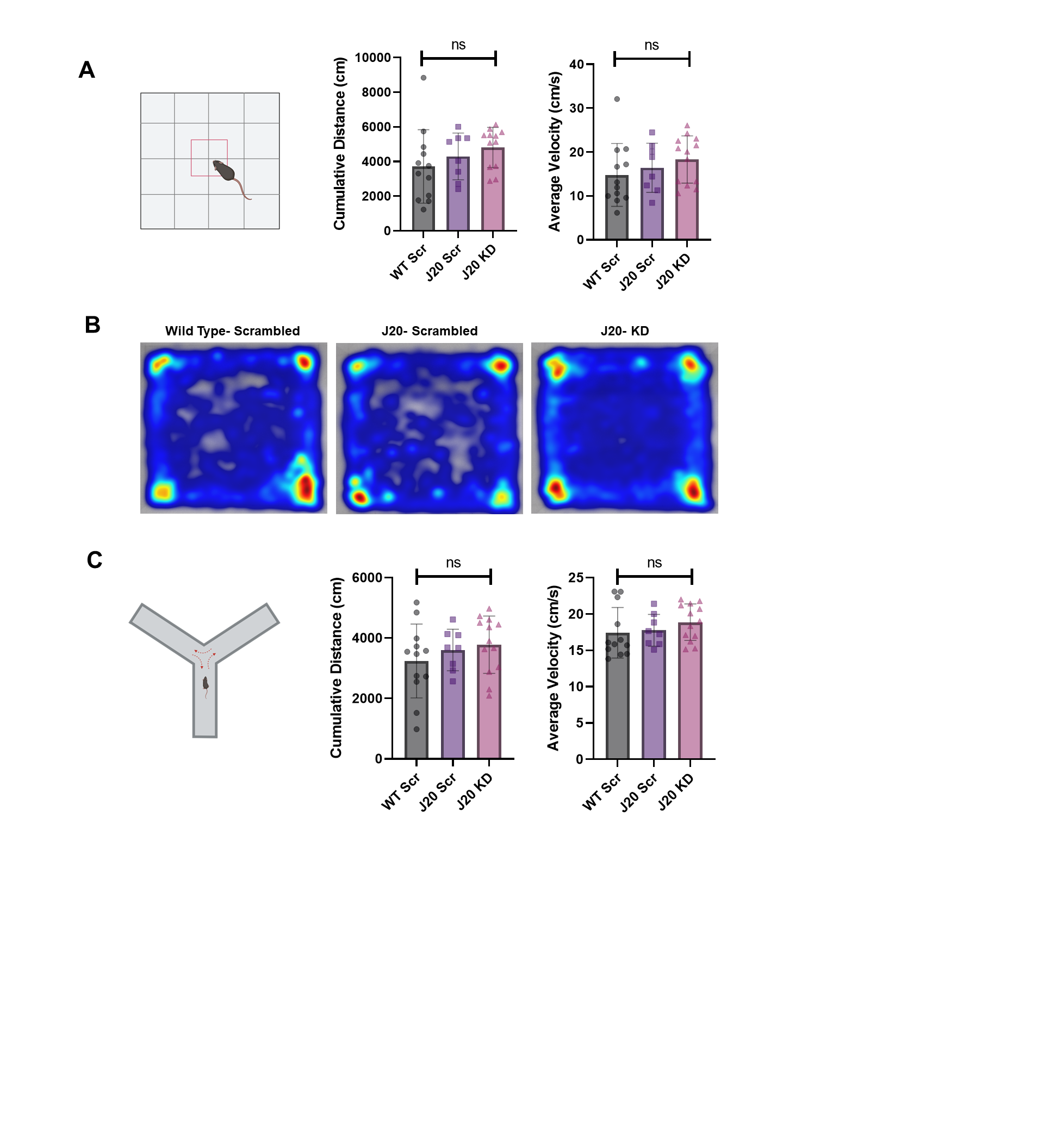
